# The Fkh1 Forkhead Associated Domain Promotes ORC Binding to a Subset of DNA Replication Origins in Budding Yeast

**DOI:** 10.1101/2021.03.01.433423

**Authors:** Timothy Hoggard, Allison J. Hollatz, Rachel Cherney, Catherine A. Fox

## Abstract

The pioneer event in eukaryotic DNA replication is binding of chromosomal DNA by the origin recognition complex (ORC), which directs the formation of origins, the specific chromosomal regions where DNA will be unwound for the initiation of DNA synthesis. In all eukaryotes, incompletely understood features of chromatin promote ORC-DNA binding. Here, we uncover a role for the Fkh1 (forkhead homolog) protein, and, in particular, its forkhead associated (FHA) domain in promoting ORC-origin binding and origin activity at a subset of origins in *Saccharomyces cerevisiae.* The majority of the FHA-dependent origins within the experimental subset examined contain a distinct Fkh1 binding site located 5’ of and proximal to their ORC sites (5’-FKH-T site). Epistasis experiments using selected FHA-dependent origins provided evidence that the FHA domain promoted origin activity through Fkh1 binding directly to this 5’ FKH-T site. Nucleotide substitutions within two of these origins that enhanced the affinity of their ORC sites for ORC bypassed these origins’ requirement for their 5’ FKH-T sites and for the FHA domain. Significantly, direct assessment of ORC-origin binding by ChIPSeq provided evidence that this mechanism affected ~25% of yeast origins. Thus, this study reveals a new mechanism to enhance ORC-origin binding in budding yeast that requires the FHA domain of the conserved cell-cycle transcription factor Fkh1.

## Introduction

Eukaryotic chromosomes rely on multiple spatially and temporally distributed DNA replication origins for their efficient and accurate duplication. Each origin has a distinct probability of activation. Thus, some origins act more efficiently and earlier in S-phase than others. These differences in origin activation probabilities generate a spatiotemporal pattern of genome duplication with relevance to genome stability and cell function [1–5]. Because chromatin heterogeneity is essential for the structure and function of chromosomes, each individual origin must act within a distinct local chromatin environment. While it is clear that differences in chromatin structure can affect the probability of origin activation, the specific features of chromatin and the molecular mechanisms by which they control origins are incompletely defined. Every yeast origin requires the same core proteins and sequential series of protein-DNA mediated reactions for its activation [6]. First, the **ORC** (**o**rigin **r**ecognition **c**omplex) selects origins by binding to the underlying chromosomal DNA [7]. In G1-phase, this ORC-DNA complex initiates a a series of molecular interactions culminating in the assembly of an inert form of the DNA replicative helicase, the MCM (**m**ini **c**hromosome **m**aintenance) complex, onto chromosomal DNA (origin licensing) [8]. The position of the MCM complex on the chromosomal DNA determines the position of the origin because, in S-phase, multiple proteins convert the loaded MCM complex into two bidirectional replicative helicases that unwind the DNA, allowing for the initiation of DNA synthesis (origin firing or activation). Given the origin-defining role of ORC-DNA interactions in this cycle, it is not surprising that chromatin’s effect on DNA accessibility plays a role in chromosomal origin distribution, with both ORC binding and origin activation being more prevalent within open, transcriptionally active chromatin regions of the genome [2, 9]. However, direct interactions between ORC and nucleosomes or non-histone chromatin-associated proteins also help recruit ORC to chromosomes, potentially promoting the formation origin-competent ORC-DNA complexes in less receptive chromatin environments [10–14].

*Saccharomyces cerevisiae* (yeast) is a powerful model for addressing the role of chromatin in controlling origins because so much is known about this organism’s ~400 origins through multiple high-resolution genome-scale studies. Data are available about ORC binding, origin efficiency (fraction of cell cycles that an origin fires in a dividing population of yeast cells) and replication time (the minutes after S-phase begins when an origin is replicated [14–21]. High-resolution data describing nucleosome occupancy and modification status as well as binding locations of relevant non-histone chromatin-associated proteins are also available [22, 23]. Because yeast ORC, unlike ORCs from other model organisms, binds to a defined DNA sequence element that is essential though not sufficient for origin function, the precise position of the origin-determining ORC-DNA interface for most yeast origins is known. Although it may seem paradoxical, the requirement for a specific, discrete ORC-DNA interface has the potential to facilitate defining chromatin-mediated mechanisms relevant to functional ORC-origin binding. In particular, while the ORC site is conserved in yeast origins, its precise sequence and affinity for ORC varies considerably between individual origins [14, 17, 24]. Thus yeast origins can be parsed by the contribution the essential ORC-DNA interface makes to their levels of ORC-origin binding *in vivo.* Origins with notably weak ORC binding sites can then serve as tools to examine how local chromatin features promote functional ORC-origin interfaces.

Previously, we identified two distinct collections of yeast origins that bind ORC using clearly contrasting mechanisms [24]. Specifically, these origins were first selected by their strong affinity for ORC *in vivo*, as defined by retention of a strong ORC ChIP signal even in *orc2-1* cells [17, 24]. Next, these origins were examined for their ORC-DNA affinity *in vitro.* Approximately 20 origins were assigned to the positive-DNA cohort (originally called DNA-dependent origins), because they contained high-affinity ORC sites (apparent Kds ranging from ~4 to 14 nM), whereas another 20 origins were assigned to the positive-chromatin cohort (originally called chromatin-dependent origins), because their ORC sites had a low affinity for ORC *in vitro* (apparent Kds ranging from ~30 nM to > 300 nM) that could not account for their strong association with ORC *in vivo.* We hypothesized that positive-chromatin origins used a feature(s) of their local chromatin environment to promote ORC-DNA binding (Figure 1).

**Figure 1.**
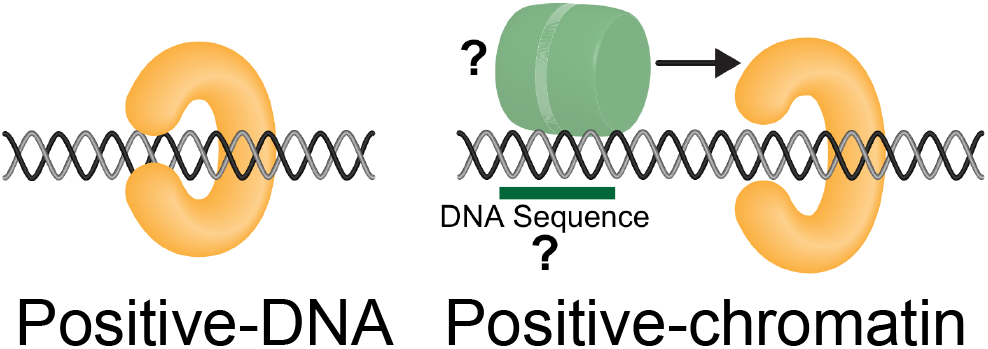
Model to explain the ORC-binding differences between two distinct yeast origin cohorts [**24**]. Origins in the positive-DNA cohort rely on direct interactions between ORC (orange crescent) and the the origin’s ORC binding site to achieve normal levels of ORC binding, while origins in the positive-chromatin cohort require additional interactions with an origin-adjacent protein(s) to achieve normal levels of ORC binding to their intrinsically weak ORC sites. The weaker ORC-DNA interactions at positive-chromatin origins have a more open ORC. In this simple model, a sequence-specific DNA binding protein (green cylinder) binds near the origin and promotes ORC binding (arrow).

Here, we applied genetic, molecular and bioinformatic approaches to compare positive-chromatin and positive-DNA origins and uncovered a role for the Fkh1 (**f**ork**h**ead **h**omolog) protein in promoting ORC-origin binding at a subset of yeast origins. The N-terminal FHA (**f**ork**h**ead **a**ssociated) domain of Fkh1, and more precisely its conserved phosphothreonine-specific protein-protein interaction function, was required by the majority (75%) of positive-chromatin origins for full activity. FHA-dependent origins were twice as likely as the FHA-independent origins to contain a FKH site in a 5’ to 3’ T-rich orientation (5’-FKH-T) positioned within 250 bp and 5’ of their ORC site. Mutational and epistasis analyses of several FHA-dependent origins indicated that this 5’ FKH-T site contributed to origin activity and accounted for these origins’ FHA-dependence. Converting the low-affinity ORC binding sites within two FHA-dependent positive-chromatin origins to high-affinity ORC sites eliminated their dependence on the Fkh1-FHA domain, supporting the conclusion that the Fkh1-FHA domain promoted ORC binding to these origins. Analyses of ORC-origin binding by ORC ChIPSeq provided evidence that FHA-dependent positive-chromatin origins and many origins outside of this strictly defined cohort required the Fkh1-FHA domain for full ORC binding *in vivo.*

## Materials and Methods

### Yeast strains and plasmids

Yeast strains were derivatives of W303, and complete genotypes are listed in supplemental Table S1. All ARS plasmids were made by cloning yeast origin fragments into the *Not1* site of pCF1897, a derivative of the pARS plasmid used in [25]. ARS sequences are listed in Table S2.

### Plasmid loss (ARS) assays

Plasmid loss assays were performed as described previously [26–28] with the exception that yeast colonies were grown in 96-well plates as described for the “tadpoling” assay [29]. Yeast transformed with an ARS were inoculated into 5 mL of selective media (lacking uracil) and grown at 25°C for ~20 hours, until cultures reached OD_600_ 1.0 - 1.5. The culture was diluted 1000-fold into 5 mL non-selective media and the diluted culture grown ~25 hours at 25°C. After each round of growth in liquid media, the culture was serially diluted into uracil-lacking or uracil-supplemented media in a 96-well plate (seven 10-fold dilutions). Yeast were grown at 25°C for 36 hours, after which colonies were counted. Plasmid loss rates (PLRs) were determined using the formula PLR = 1- (F/I)^n^ where I (Initial) is the percentage of live cells that contain a plasmid after 20 hours of selective growth, F (Final) is the percentage of live cells that contain a plasmid after ~ 25 hours of non-selective growth and n is the number of cell generations. The values presented are means of all biological replicates for the indicated ARS (≥ 3 independent yeast transformants) ± standard error.

### Basal ARS function

To determine whether *ARS1529.5* constructs in Figure 2E conferred basal ARS activity in wildtype (CFY145) or *orc2-1* (CFY266) yeast, 4 ODs of yeast were transformed with 100 ng of the indicated ARS and cells were grown on solid selective media at 23°C, the permissive growth temperature for *orc2-1* cells.

**Figure 2.**
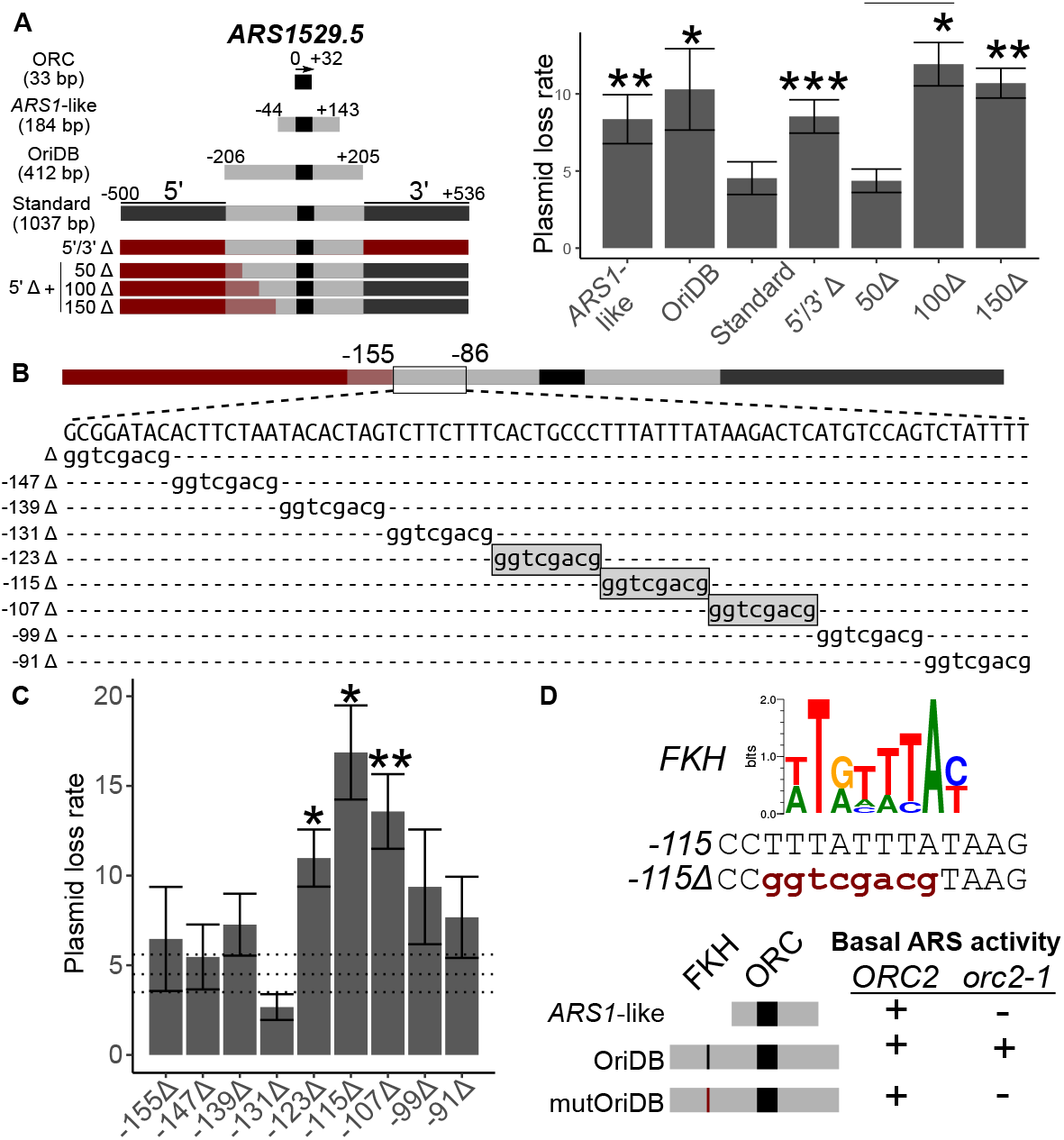
The positive-chromatin origin *ARS1529.5* required a 5’ proximal FKH binding site for optimal function. **A**) The indicated *ARS1529.5* chromosomal fragments (left) were cloned into a plasmid and the corresponding plasmid loss rates (PLRs; mean ± se) were determined (right). Asterisks here and in all subsequent figures indicate P-values for the relevant comparisons (*** < 0.001, ** < 0.01, and * < 0.05). P-values are derived from two-tailed Student’s t-tests comparing constructs to *ARS1529.5*-standard. The sequences that were used to substitute for native *ARS1529.5* sequences to create the indicated Δ clones are derived from KANMX and are denoted in red, or red shading when they overlapped OriDB regions **B)** DNA sequences between positions −155 and −86 of the *ARS1529.5* 5’Δ + 50Δ ARS construct were substituted with the 8 bp expanded *Sal*1 restriction site, gGTCGACg, as indicated. Mutations that significantly raised PLRs (P-values < 0.05) relative to *ARS1529.5*-standard are boxed in gray shading. **C)** PLRs of *Sal*1 linker-containing *ARS1529.5* mutants in B. Horizontal dashed lines represent the PLR mean ± se of the wild-type *ARS1529.5* 50Δ fragment in A. **D)** (Top) An FKH consensus motif [23]. Sequence of the wild-type region of *ARS1529.5* containing the FKH site (top), and the *Sal*I linker substitution in *ars1529.5*-FKHΔ::*Sal*1. Basal ARS activity is defined as whether colonies were recovered when yeast were transformed with the indicated ARSs. The indicated yeast cells were transformed with the *ARS1*-like version, OriDB version, or OriDB version containing the FKHΔ::*Sal*1 substitution (mutOriDB) of *ARS1529.5.* “+” indicates that colonies were recovered. “-” indicates that no colonies were recovered.

### Cell-cycle arrest and flow cytometry

To prepare genomic DNA samples for copy-number analyses, *MAT***a** cells were grown in 25 mL YPD at 30°C to an OD_600_ of ~ 0.3 at which time *α* factor was added to a concentration of 5 *μ*g/mL. Cultures continued to incubate at 30°C for 100 minutes to allow the majority of cells to reach G1-arrest. Cultures were then collected on a Whatman Nylon membrane (GE Healthcare #7404-004) using a Nalgene filtering apparatus (Thermo Scientific #300-4050) and washed with 100 mL YPD pre-warmed to 30°C. The cells were then immediately transferred to fresh pre-warmed 25 mL YPD supplemented with 200 *μ*g/mL of pronase. 2.5 mL of culture was harvested at time 0 (G1-arrested sample) and at 35 minutes (mid S-phase) after release into YPD-pronase. Genomic DNA was purified from Zymolase-treated yeast pellets using phenol-chloroform-isoamyl alcohol, followed by ethanol-based extraction. DNA was quantified on a Quantus fluorimeter. To determine cell-cycle distribution of each sample, cells were fixed with ethanol, stained with Sytox Green and analyzed by flow cytometry (BD Accuri C6 Flow Cytometer).

### Digital droplet PCR (ddPCR)

ddPCR was performed following a published protocol for examining yeast DNA replication [30] using Evagreen based chemistry (Droplet oil and Supermix; BioRad #1864005 and #1864033, respectively). ddPCR was performed on the indicated origins or non-origin control loci. Primers are listed in Table S3.

### Apparent Kds for ORC-origin binding

Recombinant yeast ORC purified from Sf9 cells was used in binding reactions with radioactively labeled origin DNA fragments as substrates as described previously [24]. The reactions were analyzed using electrophoretic mobility shift (EMSA) assays, and radioactive ORC-DNA and free DNA complexes were imaged using a GE Typhoon SLA9000 phosphoimager. Images were analyzed in ImageJ, and apparent Kd values were derived by fitting data to a one-site binding hyperbola and constraining the Bmax to 1 in Graphpad Prism 4.0. Each reaction was performed in triplicate and the mean apparent Kd was normalized to that measured for the high-affinity internal control *ARS317* (*HMR-E*) probe (appKd 7 nM).

### ORC ChIP-qPCR

ORC ChIP-qPCR was performed using the XChIP protocol [31]. 50 ml yeast cultures of three biological replicates per strain (wildtype CFY3533 and *ars1529.5-FKHΔ::Sal1* CFY4479) were grown to mid-log phase (OD_600_ ~0.5 OD) and cross-linked with 1% formaldehyde for 15 minutes. Chromatin was purified and fragmented with MNase (Worthington Biochem #LS004797) at a ratio of 0.5 MNase units/OD yeast until a 1:1 ratio of 150: 300 base pair fragments was reached (indicating a 1:1 ratio of mono- to dinucleosomes). Digested chromatin was briefly sonicated and then clarified. Ten percent of the sheared chromatin was reserved to represent starting material (Input), while the remainder was incubated with a cocktail of monoclonal antibodies against ORC as described previously [17]. Quantitative PCR was performed in a BioRad C1000 Touch thermocycler using GoTaq mix (Promega #A6001) [32]. Primer pairs for target loci are listed in Table S3. Primers were validated in a standard curve assay, demanding E and R^2^ values ≥ 0.8 and ≥ 0.95, respectively. IP and Input samples were diluted 1.2 and 2.5 fold, respectively, and technical duplicate reactions were performed. The mean of the resulting Cq values from technical duplicates were calculated, demanding technical errors ≤ 1.5% 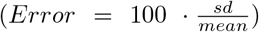. Technical means were further normalized by the relative dilution factor using the following equation: *normalized means = mean Cq* – log2(DF). Finally, percent input values were calculated for each target measured in each biological replicate for each experiment: *Percent input* = 100 · 2^(*normalized mean ip – normalized mean input*)^.

### ORC ChIPSeq

ORC ChIPSeq was performed using the XChIP protocol [31]. Crosslinked chromatin was isolated from 50 ml cultures of *FKH1* or *fkh1R80A* cells at 0.5 OD/ml. Chromatin was fragmented as described and subjected to ORC directed IP as in [14, 17]. Libraries for next generation sequencing were prepared using the Lucigen NxSeq UltraLow DNA Library Kit v2 (#15012-2). Three independent replicates of each genotype were sequenced. The ORC signal was determined for the IP and starting material and per nucleotide coverages were assessed. Coverages from like replicates were summed and then normalized for sequencing breadth and depth as described [31]. Once normalized, per nucleotide IP/input ratios were determined and mapped to 2001 bp origin fragments centered and oriented to the T-rich start of ORC site matches (n = 393). Mapped ratios from either *FKH1* or *fkh1R80A* cells at each locus were internally scaled by taking the median ratio measured within distal, flanking positions within the fragment (−1000 ~ –700 and +700 ~ +1000), and dividing each ratio within the fragment as in [31].

### Accessing genomic data

The raw data for the ORC ChIPSeq experiment can be accessed at BioProject PRJNA69402 and for the Fkh1/2 ChIPchip at GEO GSE165464.

## Results

### Nomenclature for origin fragments

The ARS assay (**a**utonomous **r**eplicating **s**equence) assay is useful for defining chromosomal DNA sequences required for yeast origin activity [25, 27, 33]. In an ARS assay, a chromosomal fragment is cloned into a bacterial plasmid backbone and plasmid replication efficiency in yeast is measured as a **p**lasmid **l**oss **r**ate (PLR). An efficient origin generates a low PLR (≤ 5% / generation). To compare positive-chromatin and positive-DNA origins in a systematic manner, we measured ARS activity of standard origin fragments from 32 origins - sixteen each from the positive-chromatin and postive-DNA cohorts. The standard fragment was defined as a 1037 bp chromosomal region centered on the origin’s ORC site (Figure 2A). The start of the ORC site’s T-rich strand was designated position ‘0’, and nucleotides 5’ and 3’ of ‘0’ were assigned negative or positive numbers, respectively. Thus the standard ARS fragment contained 500 bp 5’ and 536 bp 3’ of the ‘0’ position of the T-rich strand of the origin’s ORC site (i.e., the standard origin fragment spanned nucleotides −500 to +536). The yeast DNA replication origin database (www.OriDB.org) systematically summarizes information about each yeast chromosomal origin, including annotating the chromosomal regions that can function as ARSs as reported in the published literature [34]. For *ARS1529.5,* a positive-chromatin origin, the annotated oriDB ARS fragment is 412 bp (−206 ~ +205) and was used in some experiments. For reference, the paradigmatic yeast origin, *ARS1,* defined as the −44 to +143 chromosomal region centered on *ARS1’s* ORC site, is considerably smaller than the standard origin fragments and the *ARS1529.5-OriDB* origin fragment used here [25].

### The positive-chromatin origin *ARS1529.5* required a 5’ proximal FKH binding site for optimal activity

*ARS1529.5,* like most positive-chromatin origins, is an early and efficient origin [24]. Thus, the expectation was that the *ARS1529.5-oriDB* fragment would be an efficient ARS. However, *ARS1529.5-OriDB* and *ARS1529.5-ARS1-like* chromosomal regions produced plasmids that generated relatively high PLRs of ~10% (Figure 2B). In contrast, *ARS1529.5*-standa generated a low PLR of 5%. A construct containing a mutation of the *ARS1529.5* ORC site abolished function of *ARS1529.5*-standard, indicating that enhanced activity of this fragment was not caused by a second cryptic ORC site. The enhanced activity of *ARS1529.5*-standard was not due to insulating this ARS from inhibitory plasmid sequences because replacement of the flanking chromosomal regions with KANMX sequences reduced *ARS1529.5* activity back to that of *ARS1529.5-OriDB* (5’/3’Δ, Figure 2A).

The paradigm for yeast origin structure places key regulatory DNA elements 3’ of the T-rich strand of the essential ORC site [25]. However, a few yeast origins contain important DNA sequences 5’ of the ORC site, termed domain C, though the precise nature of these elements and their modes of action are unclear [35, 36]. Therefore, we were motivated to explore the requirement of the 5’ region of *ARS1529.5* more closely. ARS assays of additional *ARS1529.5* constructs lacking 5’ origin-adjacent sequences indicated that chromosomal DNA between nucleotides −151 and −102 contained element(s) that promoted ARS activity (Figure 2A). *Sal1* substitutions were introduced throughout the region, and ARS assays were performed on the resulting mutants (Figure 2B). Three adjacent *Sal1* substitutions reduced the function of *ARS1529.5,* with the central of these (−115Δ:*Sal*1) causing the most severe defect (Figure 2C). The −115Δ:*Sal*1 substitution mutated a sequence that matched the consensus DNA binding motif of the paralog cell-cycle transcription factors Fkh1 and Fkh2 (henceforth called an FKH motif). Thus, this result raised the possibility that either the Fkh1 or Fkh2 or both proteins bound 5’ of the T-rich strand of this origin’s ORC site to promote *ARS1529.5* activity (Figure 2D).

The screen that defined *ARS1529.5* as a positive-chromatin origin demanded that *ARS1529.5* retain ORC binding *in vivo* even under conditions where the level of functional ORC was reduced substantially by the *orc2-1* mutation [24]. If the 5’ FKH site identified above was required to promote ORC binding to *ARS1529.5*’s weak ORC site in vivo, then it should be critical for *ARS1529.5* function in *orc2-1* mutant yeast. To test this prediction, we performed a basal ARS activity assay, which simply assesses whether yeast can be transformed with a plasmid (Figure 2D). Wildtype but not *orc2-1* yeast could be transformed with the *ARS1*-like version of *ARS1529.5*. In contrast, both wildtype and *orc2-1* yeast could be transformed with *ARS1529.5-OriDB,* indicating that the additional chromosomal regions present on this fragment contained a sequence that allowed for ARS activity even when ORC function was reduced by the *orc2-1* mutation. While wildtype could be transformed with *ARS1529.5*-OriDB containing the FKH site mutation (−115Δ::*Sal*1 or *ars1529.5-FKHΔ), orc2-1* mutant yeast could not. Therefore, the 5’ FKH binding site promoted *ARS1529.5* function under conditions of limiting ORC, meeting a criterion for a postulated accessory DNA element that promotes ORC binding to positive-chromatin origins (Figure 1).

### The Fkh1-FHA domain promoted ARS activity through the FKH site positioned 5’ of the *ARS1529.5* ORC site

Fkh1 contains two conserved domains, a DNA binding domain and a **f**ork**h**ead **a**ssociated (FHA) domain (Figure 3A). FHA domains are conserved protein-binding modules with specificity for phosphorylated threonines [37, 38]. A conserved arginine (R) residue within FHA domains is essential for phospothreonine binding. A mutant allele of *FKH1, fkh1R80A,* produces normal levels of Fkh1 protein but abolishes the phosphothreonine binding function of Fkh1 [39, 40].

**Figure 3.**
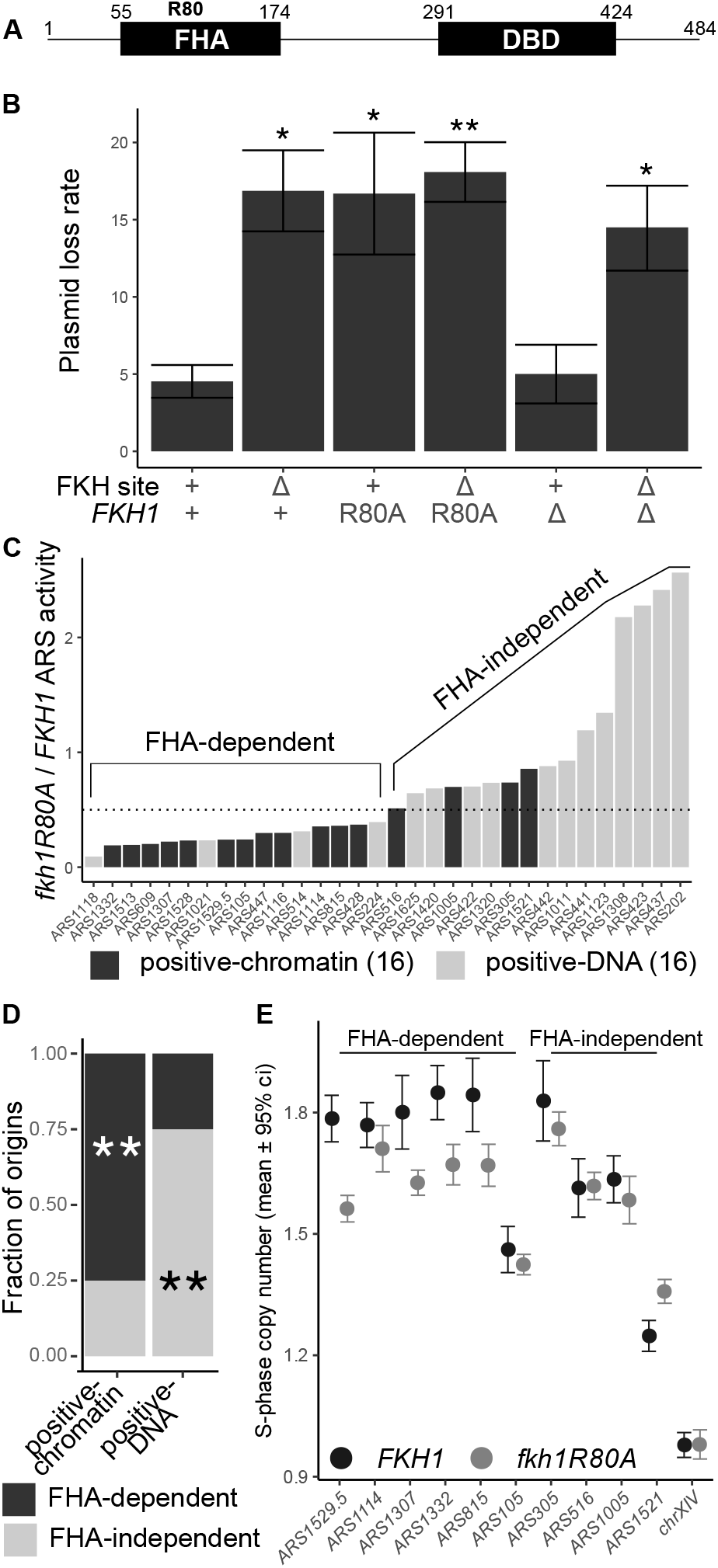
The Fkh1-FHA domain functioned through the FKH site in *ARS1529.5* and contributed to the ARS function of the majority of positive-chromatin origins. **A)** Diagram of the Fkh1 protein and relevant protein domains. Residue 80 of Fkh1 is a conserved arginine essential for the threonine-phosphopeptide-binding activity of FHA domains [44–46][38]. **B)** PLRs measured for the indicated *ARS1529.5* plasmids in *FKH1, fkh1R80A,* or *fkh1Δ* cells. **C)** ARS activities (inverse of plasmid loss rates) were determined for the indicated ARSs, cloned as standard fragments (see Figure 2A), in *fkh1R80A* and *FKH1* cells. The data are expressed as ratios of ARS activity measured in *fkh1R80A* cells to that measured in *FKH1* cells. The ARSs are ordered from most sensitive (FHA-dependent) to least affected (FHA-independent). **D)** Enrichment of FHA-dependent ARSs (*fkh1R80A*-sensitive) within the positive-chromatin and positive-DNA origin groups. P-values are derived from a hypergeometric function and significance denoted as in Fig 2. **E)** Chromosomal S-phase copy numbers as measured by ddPCR for six FHA-dependent and four FHA-independent ARSs in C. For each locus, 6 (for wild-type) or 12 (for *fkh1R80A* mutant strain) independent S-phase cell samples were assessed and the mean ± 95% confidence interval determined (see Figure S1).

To test whether the Fkh1-FHA domain contributed to *ARS1529.5* function, wild-type and *fkh1R80A* yeast were transformed with plasmids that were replicated by either wild-type *ARS1529.5* or *ars1529.5*-FKHΔ. The ARS function of wild-type *ARS1529.5* was reduced in *fkh1R80A* mutant cells to a level that was indistinguishable from that of *ars1529.5-FKHΔ* in wild-type cells, revealing that the *trans*-acting *fkh1R80A* mutation phenocopied the *cis*-acting FKH site mutation’s effect on *ARS1529.5* activity (Figure 3B). In addition, the activity of *ars1529.5-FKHΔ* was not further reduced by the *fkh1R80A* mutation, indicating that the 5’ FKH site and the Fkh1-FHA domain were epistatic. These data provided *in vivo* evidence that the Fkh1-FHA domain promoted *ARS1529.5* activity by binding to the *ARS1529.5* 5’ FKH site.

Due to their substantial sequence similarity, the paralogs Fkh1 and Fkh2 can substitute for each other’s functions in cell-cycle control of transcription [41, 42]. In addition, the majority of early-acting yeast origins that were previously defined as Fkh1/2-activated origins show altered activity only in the absence of both Fkh1 and Fkh2 [43]. However, the ARS data in Figure 3B indicated that the Fkh1-FHA domain function could explain *ARS1529.5*’s requirement for its 5’ FKH site, indicating that this defective version of Fkh1 could not be rescued by Fkh2. The simplest explanation for this outcome was that mutant fkh1R80A protein remains bound to the *ARS1529.5* 5’ FKH site. If this explanation were correct, then *fkh1Δ FKH2* cells should show no defect in *ARS1529.5* activity because, in the complete absence of Fkh1, Fkh2 should now be able to bind the 5’ FKH site. Consistent with this explanation, *fkh1Δ* cells replicated *ARS1529.5* as efficiently as wild-type cells. In addition, this ARS activity required the same 5’ FKH site that Fkh1 required (Figure 3B). Thus, in wild-type cells, the relevant *ARS1529.5* FKH site was bound by Fkh1, not Fkh2. Moreover, while the *fkh1R80A* mutant protein could bind to this site and prevent Fkh2 from binding, it could not perform a post-Fkh-binding function required for full *ARS1529.5* activity.

### The majority of positive-chromatin origins required the Fkh1-FHA domain for full activity

*ARS1529.5* was one of the twenty positive-chromatin origins identified [24]. To test whether the Fkh1-FHA domain was required by other positive-chromatin or by any positive-DNA origins, the standard versions of sixteen ARSs from each cohort were assessed for ARS activity in wild-type and *fkh1R80A* mutant cells. The PLRs for each ARS were converted to ARS activities (ARS activity = 1/PLR) and the ratio of ARS activity for each origin in *fkh1R80A* to wild-type yeast was plotted in Figure 3C. The *fkh1R80A* mutation caused a greater than two-fold reduction in ARS activity for twelve of the sixteen positive-chromatin origins. In contrast, only four of the sixteen positive-DNA ARSs showed a greater than two-fold reduction in ARS activity in *fkh1R80A* mutant yeast. Thus, while the positive-chromatin origin cohort was enriched for Fkh1FHA-dependent ARSs, the positive-DNA cohort was instead enriched for Fkh1FHA-independent ARSs (Figure 3D).

To address whether the Fkh1-FHA domain was important for the activity of FHA-dependent ARSs when they acted as origins in their native chromosomal contexts, we used droplet digital PCR (ddPCR) as a sensitive measure of DNA copy number as in [30]. For these experiments, genomic DNA was isolated from early S-phase cells after release from a G1 alpha-factor arrest (Figure S1). The early S-phase copy number of ten positive-chromatin chromosomal origins was assessed (Figure 3E); six of these showed FHA-dependent ARS activity and four did not (see Figure 3C). Four of the six FHA-dependent positive-chromatin ARSs showed significant reductions in S-phase DNA copy number in *fkh1R80A* yeast, as expected if their origin activity in their native chromosomal context was reduced [30]. In contrast, none of the FHA-independent ARSs showed replication defects. Notably, some of the individual positive-chromatin ARSs assessed here were previously identified as Fkh1/2-activated origins, yet only a subset of these showed significant replication defects in *fkh1R80A* yeast (e.g. *ARS305* is a Fkh1/2-activated origin. Yet, *ARS305* is an FHA-independent ARS (Figure 3C), and the *fkh1R80A* mutation did not affect *ARS305* chromosomal function (Figure 3E)). These data provided evidence that the FHA-dependent mechanism was distinct from the previously documented Fkh1/2-activation mechanism [23, 47, 48]. In support of this conclusion, additional bioinformatic analyses of positive-chromatin and positive-DNA origins revealed that the differences between these two origin cohorts were more tightly linked to their proposed differences in ORC-origin binding used to classify them than to either differences in their origin activation time or mode of Fkh1/2-regulation (Figure S2).

### FHA-dependent origins were enriched for T-rich-oriented FKH motifs positioned 5’ of their ORC binding sites (5’-FKH-T sites)

Previous analyses of Fkh1/2-regulated origins revealed that Fkh1/2-activated and -repressed origin groups show significant quantitative differences in Fkh1/2 binding [23]. This result is consistent with the model that Fkh1/2 act as recruitment factors to enhance the local concentration of the S-phase kinase that modifies the MCM complex and triggers origin firing [47]. However, while we could recapitulate this observation in a genome-scale Fkh1 binding experiment, the FHA-dependent and -independent origin cohorts defined in this report behaved differently, generating similar levels of Fkh1/2-association (Figure S3). Thus, while differential binding of Fkh1/2 could account for the distinct behaviors of Fkh1/2-activated and -repressed origins, this explanation was insufficient to account for the differences between FHA-dependent and FHA-independent cohorts.

To explore other possible explanations, the positions and orientations of the FKH motif matches were mapped across a 600 bp chromosomal region containing the FHA-dependent or -independent origins as defined in Figure 3C (Figure 4A-B, Figure S4). The total number of FKH motifs was similar for the FHA-dependent and -independent origin cohorts (44 motifs within the 16 FHA-dependent origin fragments versus 40 motifs among the 16 FHA-independent origin fragments). However, the two cohorts differed in how their FKH motifs were oriented and distributed relative to their ORC sites (Figure 4B-C). Specifically, origins within the FHA-dependent cohort were more likely to contain an FKH-T motif 5’ of their ORC site (5’ region; 5’-FKH-T), while origins within the FHA-independent cohort were more likely to contain an FKH-T motif 3’ to their ORC site (3’ region; 3’-FKH-T). To examine these FKH motif distributions quantitatively, the fraction of origins within the indicated cohort containing the indicated FKH motif orientation match (FKH-T or FKH-A, as in Figure 4A) was determined for the 5’ and 3’ regions of the indicated origin groups (Figure 4C). These analyses confirmed that the FHA-dependent and -independent cohorts differed from each other most strikingly in terms of the position and orientation of their FKH motifs relative to their ORC sites.

**Figure 4.**
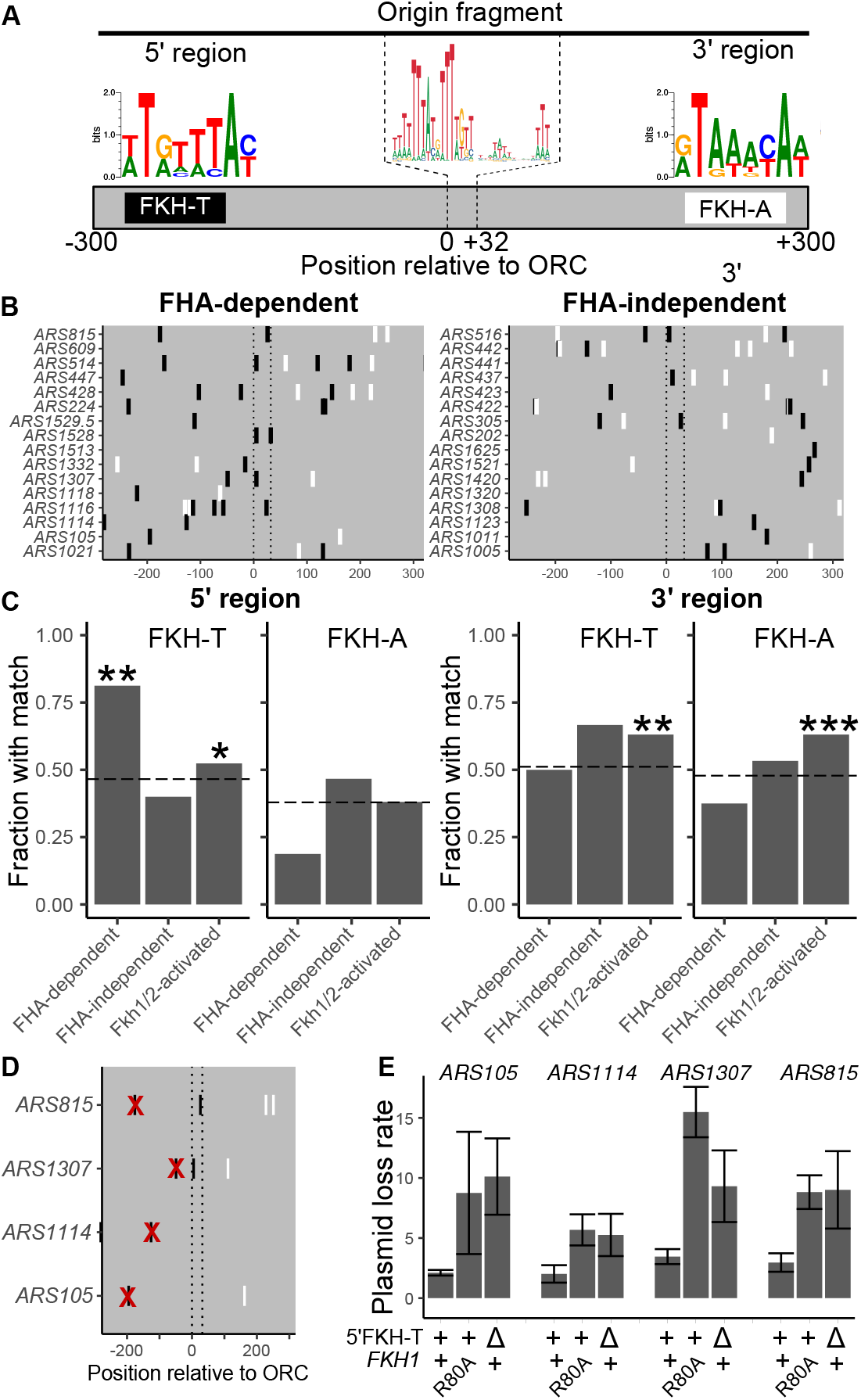
FHA-dependent ARSs were enriched for T-rich oriented FKH motifs positioned 5’ of their ORC binding sites. **A)** FHA-dependent and -independent origin fragments were examined for FKH site matches in either the T-rich (FKH-T) or the A-rich (FKH-A) orientation relative to the T-rich ORC site as depicted. Origin fragments were defined as chromosomal regions with approximately 300 nucleotides 5’ of the ORC site start of ‘0’ (−300 ~ −1, 5’ region) and 3’ of the end of the ORC site ‘+32’ (+33 —+300, 3’ region) **B)** Positions of FKH-T and FKH-A motifs in FHA-dependent (left) and FHA-independent (right) origins. Vertical dotted lines through each graph demarcate the T-rich ORC site boundaries (0 - +32) used to align each origin. **C)** Fraction of origins in the indicated cohorts (x-axis) with at least one of the indicated FKH motifs as defined in A (see also Figure S3). **D)** The 5’ FKH-T sites were replaced with an expanded *Sal1* restriction site, gGTCGACg, as with *ARS1529.5* in Figure 2. The positions of the substituted sites are indicated with a red X. E) Plasmid loss rates for the indicated *ARSs* were determined in *FKH1* or *fkh1R80A* cells.

Because the FHA-dependent origin cohort was enriched for origins with a 5’-FKH-T motif, we further examined the potential relevance of this motif. First, analyses of Fkh1/2-binding data revealed that the FHA-dependent cohort was enriched for origins that generated Fkh1-ChIP peak apexes 5’ of their ORC sites (Figure S3). Second, direct substitution of the 5’-FKH-T motif as for *ARS1529.5* in Figure 2 in four additional FHA-dependent ARSs revealed that they also required 5’-FKH-T motifs for Fkh1-FHA dependent ARS function (Figure 4E). Note that the activity of *ARS1114* was only mildly reduced by mutation of its 5’-FKH-T motif or by the *fkh1R80A* mutation, consistent with the chromosomal replication data for *ARS1114* documented in Figure 3E. Together, these observations revealed a link between a 5’-FKH-T site and FHA-dependent origin activity within the subset of origins that comprised the positive-chromatin cohort.

### A high-affinity ORC site could bypass the requirement of a 5’-FKH-T site for ARS function

The majority of positive-chromatin origins showed Fkh1-FHA-dependent ARS activity (12/16, 75%) (Figure 3C). Positive-chromatin origins were defined by the inability of their low-affinity ORC sites to explain their efficient ORC-origin association *in vivo* [24]. Therefore, we postulated that converting the low-affinity ORC sites within positive-chromatin origins to high-affinity ORC sites might bypass these origins’ requirement for the Fkh1-FHA domain for full activity (Figure 5). To address this possibility, specific nucleotide substitutions were engineered within the ORC sites of two different positive-chromatin FHA-dependent ARSs with the goal of enhancing their intrinsic affinity for ORC (defined as the affinity that ORC has for a purified origin fragment *in vitro* as measured by *in vitro* gel shift assays [24] (Figure 5A). For *ARS1529.5,* two different ORC-site-mutations were engineered: *ARS1529.5-Δ4,* where nucleotide substitutions were used to make the ORC site more similar to the consensus ORC binding motif, and *ARS1529.5-Δ514,* where the entire *ARS1529.5* ORC site (apparent Kd = 192 ± 18 nM) was replaced with the high-affinity ORC site from *ARS514* (apparent Kd = 4.1 ± 0.7 nM) [24]. For *ARS1114* (apparent Kd = 33.3 ± 2 nM), two independent ORC site mutations were generated: *ARS1114-C1T,* and *ARS1114-C30T,* with the goal of making the *ARS1114* ORC site more similar to the consensus ORC binding motif. *In vitro* ORC-DNA binding assays were performed as described previously and the binding affinity of these new ORC sites relative to a reference high-affinity ORC site (*ARS317* (*HMR-E*), apparent Kd = 7.2 ± 0.4 nM) was determined. Three of the ORC site variants, *ARS1529.5-Δ4, ARS1529.5-Δ514,* and *ARS1114-C30T* bound ORC with a high-affinity, while one variant, *ARS1114-C1T,* failed to enhance the affinity of this ORC site for ORC *in vitro* (Figure 5B).

**Figure 5.**
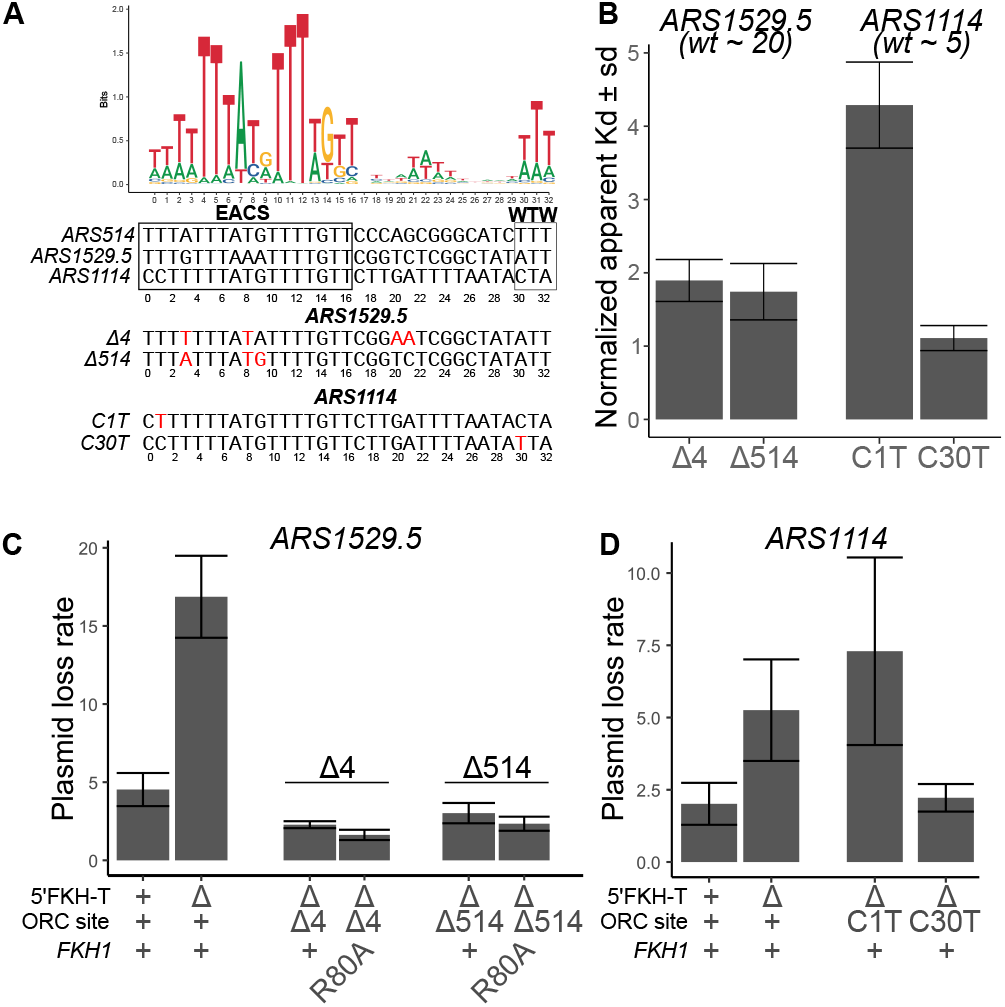
High-affinity ORC sites bypassed requirements for the 5’ FKH-T site and the FHA domain. **A)** Top: Web Logo for ORC consensus site derived from 393 chromosomal origins. Bottom: ORC binding sites for three indicated origins, and below that, the nucleotide substitutions, indicated in red, used to generate high-affinity mutant ORC binding sites. **B)** The Kds measured by gel shift assays with purified ORC and origin DNA fragments were normalized by dividing by the Kd measured for the tight-binding ORC binding site from *ARS317.* The normalized Kds for wildtype *ARS1529.5* and *ARS1114* are indicated in parentheses below the text of those origins [24]. **C)** Plasmid loss rates for wildtype and the indicated mutant versions of *ARS1529.5* in either *FKH1* or *fkh1R80A* cells. **D)** Plasmid loss rates for wild-type and the indicated mutant versions of *ARS1114* in *FKH1* cells.

The ARS activity of each of these mutant origins was determined (Figure 5C). Both of the *ARS1529.5* mutants containing high-affinity ORC sites functioned efficiently even in the absence of a 5’-FKH-T site or the absence of a functional Fkh1-FHA domain. The *ARS1114-C30T* mutant that enhanced affinity for ORC also bypassed this ARS’s dependence on its 5’ FKH-T site. However, the *ARS1114-C1T* mutation that did not enhance ORC-origin binding affinity *in vitro* still required the 5’-FKH-T site of *ARS1114* for wild-type *ARS1114* activity. Thus, a high-affinity ORC site could bypass the requirement of the 5’-FKH-T site for the efficient ARS function of these FHA-dependent positive-chromatin origins.

### The Fkh1-FHA domain promoted ORC-origin interactions at a substantial fraction of yeast chromosomal origins

The data presented above supported a specific version of the hypothesis in Figure 1 for many positive-chromatin origins. In this model, Fkh1 binds to a 5’-FKH-T binding site at these origins and uses its conserved FHA domain to promote the essential ORC-DNA interaction. To test whether the 5’-FKH-T motif promoted ORC binding to *ARS1529.5 in vivo*, chromosomal *ARS1529.5* was precisely replaced with the *ars1529.5-FKHΔ* mutant and ORC association with this locus was assessed by ORC-directed ChIP and qPCR (Figure 6A). The ORC ChIP signal generated by the *ars1529.5-FKHΔ* mutant cells was ~2.5 fold lower than the signal generated by *ARS1529.5* while the control locus was unaffected. These data provided evidence that the 5’-FKH-T motif in *ARS1529.5* identified in the experiments in Figure 2 contributed to ORC binding to *ARS1529.5* within its native chromosomal context.

**Figure 6.**
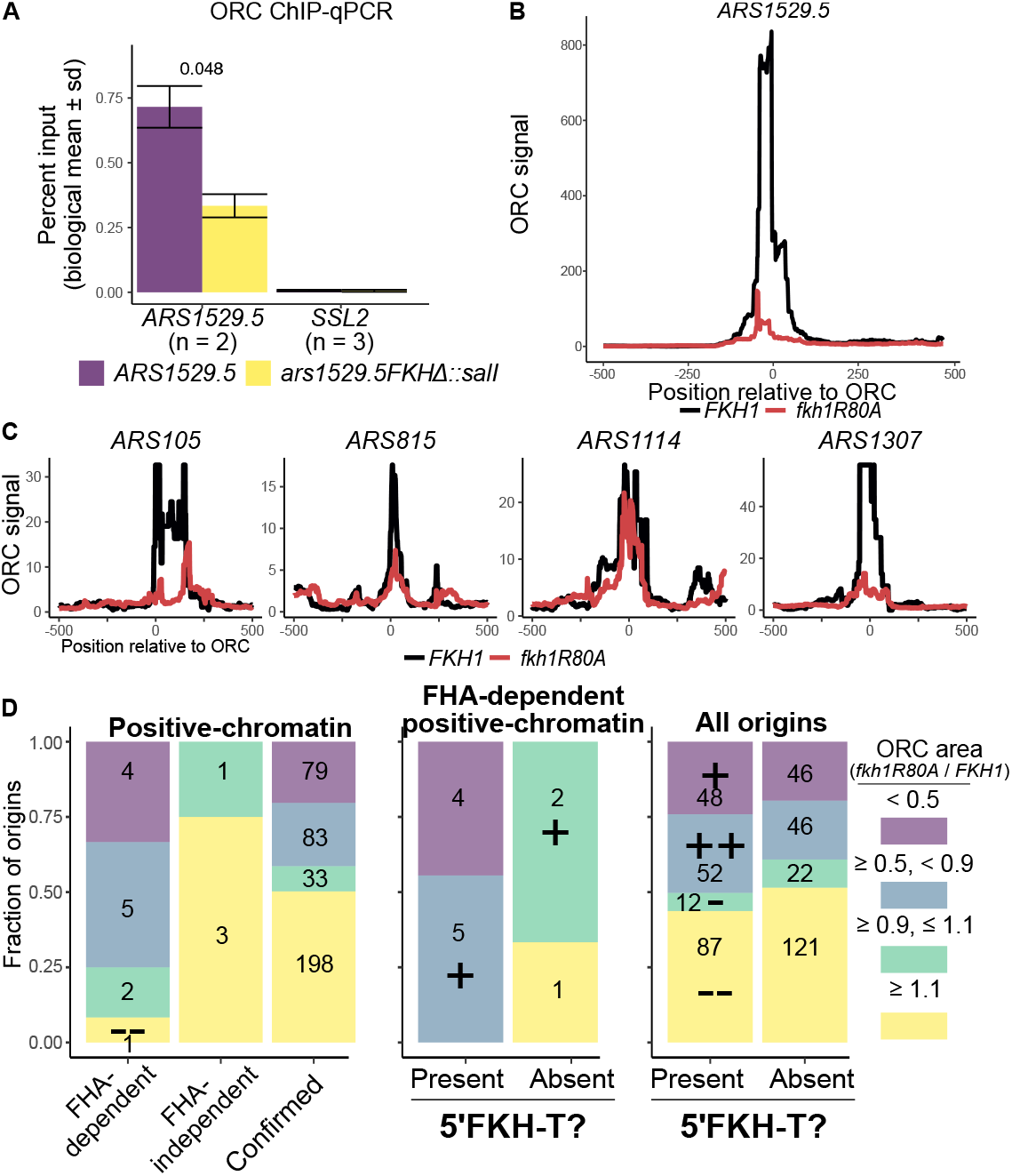
The Fkh1-FHA domain promoted ORC-origin interactions at a substantial fraction of yeast chromosomal origins. **A)** ORC signals of *ARS1529.5* and *ars1529.5-FKHΔ* in ORC ChIP-qPCR. **B)** ORC signal measured at *ARS1529.5* by ORC ChIPSeq in *FKH1* and *fkh1R80A* cells. **C)** ORC signals as in B at four additional FHA-dependent positive-chromatin origins identified in Figure 3C and with 5’ FKH matches validated in ARS assays in Figure 4D-E. **D)** Enrichment analysis of distinct groups of yeast origins defined by their ratios of *fkh1R80A*/*FKH1* ORC binding areas spanning nucleotides −100 to +100. Enrichments (indicated with a “+”) or depletion (indicated with a “–”) of categories of *fkh1R80A*/*FKH1* scaled ORC ratios were challenged against distributions of those same categories within all confirmed origins with the Hypergeometric Distribution function. ++/╌, P value < 0.01; +/−, P value < 0.05.

To further probe the roles of the Fkh1-FHA domain and a 5’-FKH-T site in ORC-origin binding, ORC ChIPSeq experiments were performed in *FKH1* and *fkh1R80A* cells. A scan of the chromosomal *ARS1529.5* locus provided evidence consistent with the locus-specific ChIP experiment in Figure 6A. Specifically, *ARS1529.5*’s association with ORC was reduced in *fkh1R80A* relative to *FKH1* cells (Figure 6B). Importantly, this lower ORC binding signal for *ARS1529.5* in *fkh1R80A* cells was not a byproduct of a non-specific genome-wide reduction in ORC-origin binding, as a majority of confirmed yeast origins (59%, 231/393) generated similar or slightly enhanced ORC signals in *fkh1R80A* cells compared to *FKH1* cells. Scans of other FHA-dependent positive-chromatin origins produced results similar to those observed for *ARS1529.5* though different quantitative effects were observed (Figure 6C). In particular, the ORC binding signal at the FHA-dependent *ARS1114* origin was only mildly reduced in *fkh1R80A* cells (Figure 6C). Interestingly, while *ARS1114* activity was reduced in *fkh1R80A* cells, it still remained a relatively efficient ARS (PLR ~ 5%) (Figure 4E) and its chromosomal replication was also not perturbed significantly (Figure 3C). These mild effects of *fkh1R80A* on *ARS1114* activity were consistent with the weak effects of this defective allele on ORC binding to this chromosomal origin.

Given the link between FHA-dependent origin activity and ORC binding revealed by the examination of several specific origins within the positive-chromatin origin cohort, we next asked whether the twelve FHA-dependent positive-chromatin origins exhibited reduced ORC binding in *fkh1R80A* cells compared to the four FHA-independent positive-chromatin origins (Figure 6D, left panel). For this analysis, we determined the ratio of the ORC binding signal in *fkh1R80A* cells to that in *FKH1* cells spanning the −100 to +100 nucleotides centered over every origin’s ORC site (n = 393). Any origin generating a 10% or greater reduction in this ratio (blue and purple) was interpreted as displaying FHA-dependent ORC binding. Every origin generating a 90% or greater *fkh1R80A*/*FKH1* ratio was interpreted as displaying FHA-independent ORC binding.

Based on these criteria, 75% (9/12) of the FHA-dependent positive-chromatin origins showed FHA-dependent ORC binding, while none of the remaining 25% (3/12) FHA-independent positive-chromatin origins did. Thus the two types of positive-chromatin origins, FHA-dependent and FHA-independent, behaved differently from each other in terms of their requirement for the Fkh1-FHA domain for ORC binding. The total number of positive-chromatin origins, n =16, was low. Thus, except for the depletion of the unaffected/slightly enhanced ORC binding category (yellow) in the Fkh1-FHA-dependent origin cohort, the P-value cut-offs for these differences relative to confirmed origins did not reach < 0.05. Therefore, to further challenge the potential relevance of this outcome further, this analysis was applied to several other distinctly defined origin groups: the positive-DNA cohort, which is comprised of origins where ORC’s affinity for the essential ORC site is sufficient to explain ORC binding *in vivo* (Figure 5, [24]) and two larger groups of Fkh1/2-regulated origins where Fkh1/2 exerts control at the S-phase MCM complex activation step, i.e. recruitment of an essential origin activation S-phase kinase [43, 47] (Figure S5). Because the positive-DNA origins, by definition, use a DNA-dependent ORC-origin binding mechanism (Figure 1), the prediction was that this origin cohort would not behave like the positive-chromatin origin cohort, and this outcome was observed (Figure S5A). Specifically, while the majority of positive-DNA origins were FHA-independent (12/16), four of them were FHA-dependent (Figure 3C). However, in contrast to the FHA-dependent positive-chromatin origins, only a small fraction (25%, 1/4) of the FHA-dependent positive-DNA origins had a reduced *fkh1R80A*/*FKH1* ORC signal. Thus, the FHA-dependent positive-DNA origins did not behave like the FHA-dependent positive-chromatin origins in terms of the effect of *fkh1R80A* on ORC binding. In addition, the Fkh1/2-regulated origins, either Fkh1/2-activated or Fkh1/2-repressed, were no more or less likely than all confirmed origins to contain origins that showed some dependence on the Fkh1-FHA domain for ORC binding (Figure S5B). Together these analyses supported the conclusion that the FHA-dependent positive-chromatin cohort was distinct in its enrichment for origins that used an Fkh1-FHA-mechanism to promote ORC binding.

Next, the FHA-dependent positive-chromatin origins were further analyzed to test the relationship between a 5’-FKH-T site and Fkh1-FHA dependent ORC binding. Specifically, even within the FHA-dependent positive-chromatin subcohort (n=12), while the majority (75%, 9/12) showed FHA-dependent ORC binding, a minority did not (25%, 3/12). Therefore, we further parsed the origins within this subcohort by whether they contained a 5’-FKH-T motif (Figure 6D, middle panel). Strikingly, 100% of FHA-dependent positive-chromatin origins with a 5’-FKH-T motif (n = 9) had FHA-dependent ORC binding. In contrast, none of the three remaining FHA-dependent positive-chromatin origins that lacked a 5’-FKH-T motif showed FHA-dependent ORC binding. Thus, Fkh1 binding to a 5’-FKH-T site was the dominant mechanism that predicted FHA-dependent ORC binding among positive-chromatin origins. Nevertheless, even within this stringently defined origin cohort, FHA-dependent ARS activity was not obligatorily linked to FHA-dependent ORC-origin binding in a native chromosomal context.

While Fkh1-FHA-dependent ORC binding was a predominant mechanism within the positive-chromatin cohort, this cohort was intentionally restricted to a small number of yeast origins (~5%) that met arbitrary stringent cut-offs for ORC binding strengths *in vivo* and *in vitro* [24]. However, it was clear that many more confirmed origins showed FHA-dependent reductions in ORC binding signals; 42% (162/393) of confirmed origins fell into the FHA-dependent ORC binding category (Figure 6D, left panel). To address whether a link to a 5’-FKH-T site and FHA-dependent ORC binding could be revealed by examining all confirmed origins, the confirmed origins were parsed by the presence of a 5’-FKH-T site (Figure 6D, right panel). Notably, the fraction of confirmed origins that contained at least one 5’-FKH-T motif was significantly enriched for origins that showed FHA-dependent ORC binding. These observations suggested that the mechanism identified at the majority of FHA-dependent positive-chromatin origins might operate at as many as 25% (100/393) of confirmed yeast origins.

## Discussion

Budding yeast ORC binds to a specific DNA element within yeast origins that is essential for origin activity, a distinct feature of yeast origins that has made this organism so useful for identifying the core origin-binding proteins in eukaryotic cells, including ORC [6]. However, while sequence-specific binding by ORC is important for defining yeast origins, several lines of evidence indicate that yeast, like other eukaryotes, also uses incompletely defined features of origin-adjacent chromatin to promote ORC-origin binding [14, 16, 24, 49, 50]. The experiments in this study revealed that Fkh1 was an origin-adjacent chromatin-associated protein that promotes ORC-origin binding and origin activity at a subset of origins. This Fkh1-dependent mechanism required the conserved Fkh1-FHA domain and a distinct Fkh1 binding site located 5’ of the essential ORC site. Thus, stabilization of the essential ORC-DNA interaction is a mechanism by which Fkh1 promotes the formation of chromosomal origins in budding yeast.

### Fkh1 promotes origin activity at multiple steps in the origin cycle

Fkh1 and its paralog Fkh2 positively regulate ~20% of yeast chromosomal origins, and the majority of these Fkh1/2-activated origins (n = 84) are among the earliest replicating origins in this organism [43, 51]. Several lines of evidence indicated that the Fkh1-FHA-dependent mechanism described here was distinct from the mechanism operating at Fkh1/2-activated origins. In particular, Fkh1/2-activated origins are regulated primarily at the S-phase activation step by Fkh1/2 proteins adjacently bound to origins and recruiting through direct interactions the limiting S-phase kinase, DDK (**D**bf4-**d**ependent **k**inase)[47, 48]. In contrast, the Fkh1-FHA-dependent mechanism could be explained by enhanced binding of ORC to origin DNA. Evidence supporting this conclusion included direct assessment of ORC-origin binding *in vivo* and the conversion of FHA-dependent positive-chromatin ARSs to FHA-independent ARSs by engineering mutant ORC sites with enhanced affinities for ORC. In addition, the FHA-dependent mechanism was more tightly linked to ORC-origin binding mechanisms (e.g. positive-chromatin compared to the control/contrast collection of positive-DNA origins) than to either origin activation time or modes of Fkh1/2-regulation. The Fkh1/2-activated mechanism is achieved through a high local concentration of origin-adjacent FKH sites that bind Fkh1/2 that in turn establish a high local concentration of the limiting DDK that triggers origin firing. Consistent with this model, Fkh1/2-activated origins show significantly greater association with Fkh1/2 proteins than Fkh1/2-repressed origins do, accounting for their enhanced competitiveness for the DDK [23]. In contrast, FHA-dependent and FHA-independent origins contained a similar number of origin-adjacent FKH motifs and showed similar levels of association with Fkh1, providing evidence that a quantitative difference in Fkh1 binding could not easily account for differences in these origins’ dependencies on the Fkh1-FHA domain. However, while the Fkh1/2-activated and Fkh1-FHA-dependent mechanisms represent distinct mechanisms for how Fkh1 is used to enhance origin activity, these mechanisms are not mutually exclusive. Indeed, while *ARS305* is an Fkh1/2-activated but Fkh1-FHA-independent origin, *ARS1529.5* is both Fkh1/2-activated and Fkh1-FHA-dependent. Thus, Fkh1 bound to *ARS1529.5* may both stabilize ORC and help recruit the DDK, whereas at other origins it may perform only one of these functions.

### A distinct structural class of yeast origins: The Fkh1-FHA domain acted through an FKH site positioned 5’ of the essential origin ORC site

An ORC site is an essential but not a sufficient element for yeast origin activity. An influential paradigm for yeast origin structure positions the accessory elements required for normal levels of origin activity 3’ of the T-rich strand of the essential ORC site [25]. These 3’ accessory elements are AT-rich and often include near matches to the ORC binding site in the A-rich orientation (i.e. orientation opposite to that of the essential ORC site). Because the Fkh1/2 core binding motif is also AT-rich, these 3’ accessory elements also often contain matches to FKH motifs. The potentially overlapping biochemical functions of these 3’ accessory elements can make assigning their definitive roles at origins challenging. For example, molecular and biochemical studies provide evidence that a match to a reverse ORC site can enhance binding of a second ORC that aids in the loading of the second MCM hexamer during the formation of the double-hexamer MCM complex [52, 53]. Mutational analyses of a few Fkh1/2-activated origins indicate that FKH motifs positioned 3’ of the essential T-rich ORC site are required for their activation, consistent with these sites acting as Fkh1/2 binding sites *in vivo* [43, 54]. However, it is difficult to discern whether mutations of these FKH motifs reduce origin activity because they abolish Fkh1/2 binding or because they reduce the binding of a second ORC or both. Either effect could conceivably alter the sensitivity of the origin to Fkh1/2 activation.

In this report, a definitive role for the 5’-FKH-T motif in several Fkh1-FHA-dependent positive-chromatin origins could be assigned because the *fkh1R80A* allele precisely abolished the established function of the FHA domain in phosphothreonine peptide binding while leaving Fkh1’s DNA binding domain intact [39, 40, 46]. Thus, epistasis experiments generated compelling evidence that the Fkh1-FHA domain acted directly through these origins’ 5’ FKH sites. Specfically, the activity of a model Fkh1-FHA-dependent origin, *ARS1529.5*, was reduced substantially and to the same extent in *fkh1R80A* or in *FKH1* cells containing a substitution of the 5’-FKH-T site in *ARS1529.5.* Moreover, yeast containing both mutations, a *fkh1R80A* allele and the 5’-FKH-T site substitution, did not reduce the activity of *ARS1529.5* further than either mutation alone. Consistent results were observed with other Fkh1-FHA-dependent origins. While studies of a small number of origins indicate that some yeast origins contain element(s) 5’ of their ORC sites that contribute to origin activity, a role for these elements has not been assigned [35, 36]. The data presented here provide evidence that a FKH site can be an important 5’ origin-accessory element in yeast.

While experimental data provided evidence that the majority (56%, 9/16) of positive-chromatin origins used the 5’-FKH-T, Fkh1-FHA-dependent mechanism reported here, these origins represent only about two percent of confirmed yeast origins. However, the genome-scale analyses of ORC binding provided evidence that the Fkh1-FHA domain was relevant to competitive levels of ORC binding at ~40% origins, suggesting this domain might be used broadly across the genome to enhance ORC-origin selection. Moreover, a significant link between an origin-adjacent 5’-FKH-T motif and Fkh1-FHA-dependent ORC binding was revealed by analyses of all 393 confirmed origins, suggesting that the less familiar origin structure and mechanism uncovered by a study of positive-chromatin origins could operate more generally, affecting 25% of yeast origins. Notably, this value may be an underestimate, as FKH motif searches were confined to 250 bp 5’ of the ORC site. Thus, taken together with what is known about the mechanism that controls Fkh1/2-activated origins, this study reveals that Fkh1, and possibly Fkh2, may contribute directly to origin regulation by influencing several distinct steps of the origin cycle in both G1- and S-phase. A clearer picture of the roles for Fkh proteins in origin control and the relationship of this control to other genomic structures and processes will require higher-resolution molecular information about Fkh1-chromosome and Fkh1-protein interactions.

## Acknowledgments

We are grateful for the multiple undergraduate researchers who participated in the Team Origin Undergraduate Research Project, notably Christen Geyer, Cassaundra Burt, Yanzi Jiang, and Troy Meikle, and helped clone ARSs and performed ARS assays that influenced the experiments used in this report. We are grateful to Laura Vanderploeg (MediaLab, UW-Madison Biochemistry Department) for Figure 1. We also thank Mike Lodes and Rob Brazas (Lucigen Corporation, Madison, WI) for generating and sequencing the ORC ChIP libraries, and Xiaolan Zhao (Memorial Sloan Kettering Cancer Center) and Erika Shor (Center for Discovery and Innovation, Hackensack Meridian Health) for helpful suggestions on the manuscript.

## Funding

This study was supported by an NIH R01 grant (NIGMS 056890 - CAF) and a Teaching and Learning Innovation Award (Team Origin Undergraduate Research Project) from UW Madison.

## Notes

### Competing Interest Statement

The authors have declared no competing interest.

## References

1. Debatisse, M., Le, T. B., Letessier, A., Dutrillaux, B. & Brison, O. Common fragile sites: mechanisms of instability revisited. Trends Genet 28, 22–32 (Jan. 2012).

2. Miotto, B., Ji, Z. & Struhl, K. Selectivity of ORC binding sites and the relation to replication timing, fragile sites, and deletions in cancers. Proc Natl Acad Sci U S A 113, E4810–9 (Aug. 2016).

3. van, B. A., Buchanan, C., Charboneau, E., Fangman, W. & Brewer, B. An origin-deficient yeast artificial chromosome triggers a cell cycle checkpoint. Mol Cell 7, 705–13 (Apr. 2001).

4. Müller, C. & Nieduszynski, C. DNA replication timing influences gene expression level. J Cell Biol 216, 1907–1914 (July 2017).

5. Rivera-Mulia, J. et al. Replication timing networks reveal a link between transcription regulatory circuits and replication timing control. Genome Res 29, 1415–1428 (Sept. 2019).

6. Bell, S. & Labib, K. Chromosome Duplication in Saccharomyces cerevisiae. Genetics 203, 1027–67 (July 2016).

7. Bell, S. & Stillman, B. ATP-dependent recognition of eukaryotic origins of DNA replication by a multiprotein complex. Nature 357, 128–34 (May 1992).

8. Remus, D. & Diffley, J. Eukaryotic DNA replication control: lock and load, then fire. Curr Opin Cell Biol 21, 771–7 (Dec. 2009).

9. Lubelsky, Y. et al. DNA replication and transcription programs respond to the same chromatin cues. Genome Res 24, 1102–14 (July 2014).

10. Thomae, A. et al. Interaction between HMGA1a and the origin recognition complex creates site-specific replication origins. Proc Natl Acad Sci U S A 105, 1692–7 (Feb. 2008).

11. Norseen, J. et al. RNA-dependent recruitment of the origin recognition complex. EMBO J 27, 3024–35 (Nov. 2008).

12. Hayashi, M. & Masukata, H. Regulation of DNA replication by chromatin structures: accessibility and recruitment. Chromosoma 120, 39–46 (Feb. 2011).

13. Shen, Z. et al. A WD-repeat protein stabilizes ORC binding to chromatin. Mol Cell 40, 99–111 (Oct. 2010).

14. Müller, P. et al. The conserved bromo-adjacent homology domain of yeast Orc1 functions in the selection of DNA replication origins within chromatin. Genes Dev 24, 1418–33 (July 2010).

15. Wyrick, J. et al. Genome-wide distribution of ORC and MCM proteins in S. cerevisiae: high-resolution mapping of replication origins. Science 294, 2357–60 (Dec. 2001).

16. Eaton, M., Galani, K., Kang, S., Bell, S. & MacAlpine, D. Conserved nucleosome positioning defines replication origins. Genes Dev 24, 748–53 (Apr. 2010).

17. Shor, E. et al. The origin recognition complex interacts with a subset of metabolic genes tightly linked to origins of replication. PLoS Genet 5, e1000755 (Dec. 2009).

18. McGuffee, S., Smith, D. & Whitehouse, I. Quantitative, genome-wide analysis of eukaryotic replication initiation and termination. Mol Cell 50, 123–35 (Apr. 2013).

19. Yabuki, N., Terashima, H. & Kitada, K. Mapping of early firing origins on a replication profile of budding yeast. Genes Cells 7, 781–9 (Aug. 2002).

20. Raghuraman, M. et al. Replication dynamics of the yeast genome. Science 294, 115–21 (Oct. 2001).

21. Feng, W. et al. Genomic mapping of single-stranded DNA in hydroxyurea-challenged yeasts identifies origins of replication. Nat Cell Biol 8, 148–55 (Feb. 2006).

22. Weiner, A. et al. High-resolution chromatin dynamics during a yeast stress response. Mol Cell 58, 371–86 (Apr. 2015).

23. Ostrow, A. et al. Fkh1 and Fkh2 bind multiple chromosomal elements in the S. cerevisiae genome with distinct specificities and cell cycle dynamics. PLoS One 9, e87647 (2014).

24. Hoggard, T., Shor, E., Müller, C., Nieduszynski, C. & Fox, C. A Link between ORC-origin binding mechanisms and origin activation time revealed in budding yeast. PLoS Genet 9, e1003798 (2013).

25. Marahrens, Y. & Stillman, B. A yeast chromosomal origin of DNA replication defined by multiple functional elements. Science 255, 817–23 (Feb. 1992).

26. Palzkill, T., Oliver, S. & Newlon, C. DNA sequence analysis of ARS elements from chromosome III of Saccharomyces cerevisiae: identification of a new conserved sequence. Nucleic Acids Res 14, 6247–64 (Aug. 1986).

27. Chang, F. et al. High-resolution analysis of four efficient yeast replication origins reveals new insights into the ORC and putative MCM binding elements. Nucleic Acids Res 39, 6523–35 (Aug. 2011).

28. Hoggard, T. et al. High Throughput Analyses of Budding Yeast ARSs Reveal New DNA Elements Capable of Conferring Centromere-Independent Plasmid Propagation. G3 (Bethesda) 6, 993–1012 (Apr. 2016).

29. Welch, A. & Koshland, D. A simple colony-formation assay in liquid medium, termed ‘tadpoling’, provides a sensitive measure of Saccharomyces cerevisiae culture viability. Yeast 30, 501–9 (Dec. 2013).

30. Batrakou, D., Heron, E. & Nieduszynski, C. Rapid high-resolution measurement of DNA replication timing by droplet digital PCR. Nucleic Acids Res 46, e112 (Nov. 2018).

31. Skene, P. & Henikoff, S. A simple method for generating high-resolution maps of genome-wide protein binding. Elife 4, e09225 (June 2015).

32. Radman-Livaja, M. et al. Dynamics of Sir3 spreading in budding yeast: secondary recruitment sites and euchromatic localization. EMBO J 30, 1012–26 (Mar. 2011).

33. Palzkill, T. & Newlon, C. A yeast replication origin consists of multiple copies of a small conserved sequence. Cell 53, 441–50 (May 1988).

34. Siow, C., Nieduszynska, S., Müller, C. & Nieduszynski, C. OriDB, the DNA replication origin database updated and extended. Nucleic Acids Res 40, D682–6 (Jan. 2012).

35. Newlon, C. & Theis, J. The structure and function of yeast ARS elements. Curr Opin Genet Dev 3, 752–8 (Oct. 1993).

36. Raychaudhuri, S., Byers, R., Upton, T. & Eisenberg, S. Functional analysis of a replication origin from Saccharomyces cerevisiae: identification of a new replication enhancer. Nucleic Acids Res 25, 5057–64 (Dec. 1997).

37. Durocher, D., Henckel, J., Fersht, A. & Jackson, S. The FHA domain is a modular phosphopeptide recognition motif. Mol Cell 4, 387–94 (Sept. 1999).

38. Reinhardt, H. & Yaffe, M. Phospho-Ser/Thr-binding domains: navigating the cell cycle and DNA damage response. Nat Rev Mol Cell Biol 14, 563–80 (Sept. 2013).

39. Dummer, A. et al. Binding of the Fkh1 Forkhead Associated Domain to a Phosphopeptide within the Mph1 DNA Helicase Regulates Mating-Type Switching in Budding Yeast. PLoS Genet 12, e1006094 (June 2016).

40. Li, J. et al. Regulation of budding yeast mating-type switching donor preference by the FHA domain of Fkh1. PLoS Genet 8, e1002630 (2012).

41. Hollenhorst, P., Bose, M., Mielke, M., Müller, U. & Fox, C. Forkhead genes in transcriptional silencing, cell morphology and the cell cycle. Overlapping and distinct functions for FKH1 and FKH2 in Saccharomyces cerevisiae. Genetics 154, 1533–48 (Apr. 2000).

42. Hollenhorst, P., Pietz, G. & Fox, C. Mechanisms controlling differential promoter-occupancy by the yeast forkhead proteins Fkh1p and Fkh2p: implications for regulating the cell cycle and differentiation. Genes Dev 15, 2445–56 (Sept. 2001).

43. Knott, S. et al. Forkhead transcription factors establish origin timing and long-range clustering in S. cerevisiae. Cell 148, 99–111 (Jan. 2012).

44. Durocher, D., Smerdon, S., Yaffe, M. & Jackson, S. The FHA domain in DNA repair and checkpoint signaling. Cold Spring Harb Symp Quant Biol 65, 423–31 (2000).

45. Durocher, D. & Jackson, S. The FHA domain. FEBS Lett 513, 58–66 (Feb. 2002).

46. Durocher, D. et al. The molecular basis of FHA domain:phosphopeptide binding specificity and implications for phospho-dependent signaling mechanisms. Mol Cell 6, 1169–82 (Nov. 2000).

47. Fang, D. et al. Dbf4 recruitment by forkhead transcription factors defines an upstream rate-limiting step in determining origin firing timing. Genes Dev 31, 2405–2415 (Dec. 2017).

48. Zhang, H. et al. Dynamic relocalization of replication origins by Fkh1 requires execution of DDK function and Cdc45 loading at origins. Elife 8 (May 2019).

49. Hu, Y. et al. Evolution of DNA replication origin specification and gene silencing mechanisms. Nat Commun 11, 5175 (Oct. 2020).

50. Lee, C. et al. Humanizing the yeast origin recognition complex. Nat Commun 12, 33 (Jan. 2021).

51. Aparicio, O. Location, location, location: it’s all in the timing for replication origins. Genes Dev 27, 117–28 (Jan. 2013).

52. Coster, G. & Diffley, J. Bidirectional eukaryotic DNA replication is established by quasi-symmetrical helicase loading. Science 357, 314–318 (July 2017).

53. Miller, T., Locke, J., Greiwe, J., Diffley, J. & Costa, A. Mechanism of head-to-head MCM double-hexamer formation revealed by cryo-EM. Nature 575, 704–710 (Nov. 2019).

54. Reinapae, A. et al. Recruitment of Fkh1 to replication origins requires precisely positioned Fkh1/2 binding sites and concurrent assembly of the pre-replicative complex. PLoS Genet 13, e1006588 (Jan. 2017).

