## Supplemental methods and figures for "The Fkh1 Forkhead Associated Domain Promotes ORC Binding to a Subset of DNA Replication Origins in Budding Yeast"

#### Fkh1 ChIPchip

The Fkh1 ChIP experiment was performed on two independent isolates each from congenic strains, *FKH1* and *fkh1*::*HISG*, and hybridized to a high density tiled array as described in [1]. Fkh1-associated fragments were enriched using polyclonal antibody raised against the N-terminal region of Fkh1 [2, 3]. For each strain the log2 ratios for each probe of the two experiments were averaged. To control for potential cross reactivity of the polyclonal antibody for Fkh2, the log2 ratios from each array tile from the *FKH1* experiment were normalized by dividing values by those measured in the *fkh1* experiment. To map normalized Fkh1 ChIP signals at origins, 1201 bp origin-containing fragments were aligned relative to the T-rich start of their ORC site match, position 0, such that 600 nucleotides 5' of the ORC site start (negative numbers) and 600 nucleotides 3' of the ORC site (positive numbers) were assessed. The log2 ratios of ChIP/input were mapped to these fragments to give unscaled Fkh1 ChIP ratios for each locus analyzed. To compare Fkh1 binding between different origin cohorts within a given strain, the Fkh1 signal at each nucleotide within each fragment analyzed was scaled by the median log2 ratio measured between position -1200 and -1000 (interpreted as "background" signal for that region) [4]. We then determined the distribution of scaled Fkh1 ChIP ratios at each nucleotide for each fragment within an indicated origin cohort and plotted the median Fkh1 ChIP signal for the indicated group of origins including quartiles 1 and 3 for the variation within the collection.

#### Deriving and challenging FKH consensus motif from the Fkh1/2 ChIPchip

The program ChIPOTle (parameters of 400 and 100 bp window and step size, respectively) was used to identify peaks meeting -log10 P-value 3 [5]. The motif finder tool in MochiView [6] was used to identify motifs enriched within Fkh1 peaks, specifying an 8 bp motif and using a 4th order Hidden Markov Model of the entire genome. The motif identified agreed with that previously observed. To validate the FKH motif consensus in this study, the R package DiffLogo [7] was used to compare this study's consensus with that previously identified.

### Supplemental Figures

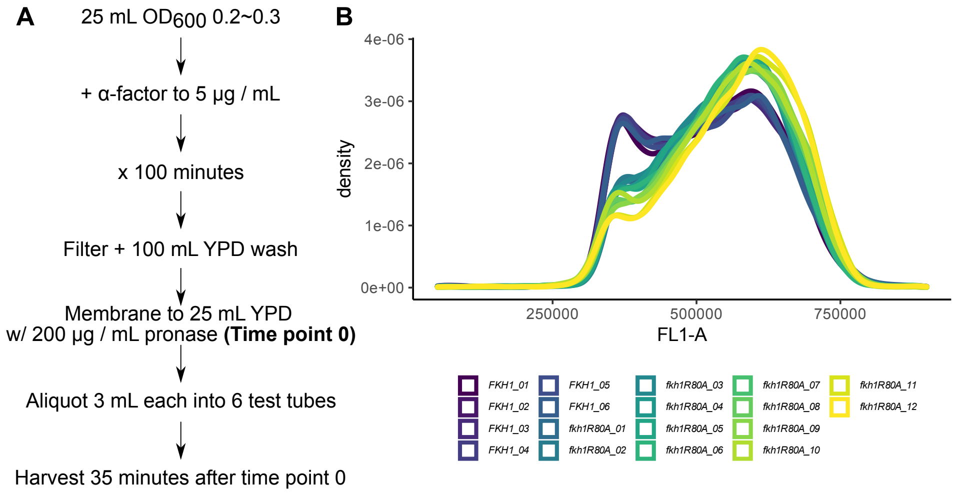


**Figure S1.** Determining replication behavior of positive-chromatin origins in their native chromosomal locations. In Figure 3F, ddPCR was used to assess the average S-phase copy number of selected origin loci from six cultures of *FKH1* and twelve cultures of *fkh1R80A* cells that had been released from G1-arrest. **A)** Schematic of arrest and release protocol used to obtain the data for Figure 3F. **B)** Flow Cytometry of six FKH1 and twelve *fkh1R80A* cells at time of harvest for ddPCR.

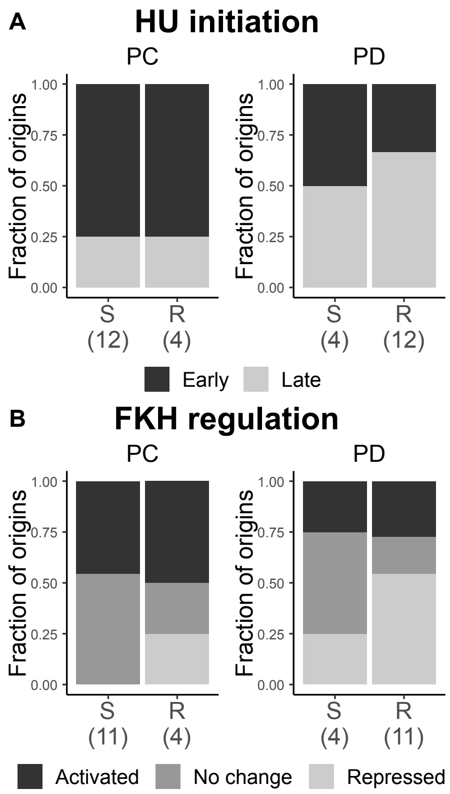


**Figure S2.** Relationship between Fkh1-FHA-regulation and origin-activation time or mode of Fkh1/2-regulation. Analyses were performed to ask whether the effect of the *fkh1R80A* mutation on the activity the 32 ARSs in Figure 3C was linked significantly to differences between their replication times as chromosomal origins (i.e. early versus late activated, as defined by hydroxyurea (HU) resistant or sensitive origin activity respectively [8, 9] or previously defined Fkh1/2 regulation [10]. PC refers to the cohort of 16 positive-chromatin origins and PD to the 16 positive-DNA origins assessed in this study in Figure 3C. S refers to the origins in Figure 3C whose ARS activity was reduced $\geq$ 2-fold in *fkh1R80A* compared to *FKH1* cells, while R refers the origins in Figure 3C whose ARS activity was reduced $<$ 2-fold in *fkh1R80A* compared to *FKH1* cells. HU Initiation refers to the method used to classify Early (HU resistant) and Late (HU sensitive) origins. Early origins are active (fire) in cells released from G1 arrest into S-phase in the presence of 200 mM HU, while Late origins are not (do not fire) under these conditions. Fkh1/2 regulation refers to origins that were classified as dependent on Fkh1/2 for firing (Activated), origins that were unaffected by Fkh1/2 (No change) and origins that were repressed by Fkh1/2 (Repressed) as in [10]. **A)** The 32 origins in Figure 3C (16 positive-chromatin and 16 positive-DNA origins) were parsed into two groups, Fkh1-FHA-dependent (S) and Fkh1-FHA-independent (R) and the fraction of Early and Late Origins within each sub-category was determined. **B)** The same sub-categories of origins assessed in A were examined for the fraction of origins that were assigned the indicated type of Fkh1/2 regulation (Activated, No Change or Repressed) as in [10].

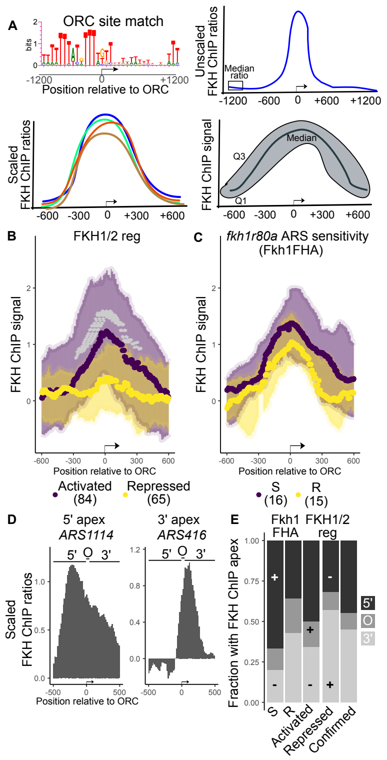


**Figure S3.** **A)** Schematic of the Fkh1/2 ChIPchip. **B)** The Fkh1 per-nucleotide median ChIP signals at Fkh1/2-activated (purple) and Fkh1/2-repressed (yellow) origins as defined in [10] were plotted as outlined in A. The grey * marks nucleotides whose differences in Fkh1 binding signals between the two groups of origins met a P-value of $< 0.05$ (Wilcoxon Rank Sum Test). The number of origins in each of cohorts is indicated in parenthesis at the bottom of each panel. **C)** The same analyses was applied to the smaller cohorts of FHA-dependent (*fkh1R80A*-sensitive, S) and FHA-independent (*fkh1R80A*-resistant, R) as defined in Figure 3C of this report. While the FHA-independent origins (yellow) showed a lower median value for Fkh1 binding signals than the FHA-dependent origins (purple), no difference at any nuleotide position along these aligned loci met a P-value cut-of $< 0.05$ ). **D)** The position of the apex of the Fkh1 ChIP signals were mapped at each origin and assigned a 5' position relative to the ORC site (as shown for *ARS1114*), directly over the ORC site (O) position, or a position 3' to the ORC site (as shown for *ARS1 - ARS416*). **E)** The fraction of origins showing the given type of Fkh1 apex (y-axis) was determined for each indicated origin cohort (x-axis). S and R indicate FHA-dependent and FHA-independent origin cohorts defined in this report in Figure 3C, respectively. The Activated and Repressed origin cohorts refer to the Fkh1/2-activated and Fkh1/2-repressed origins, respectively, defined in [10].

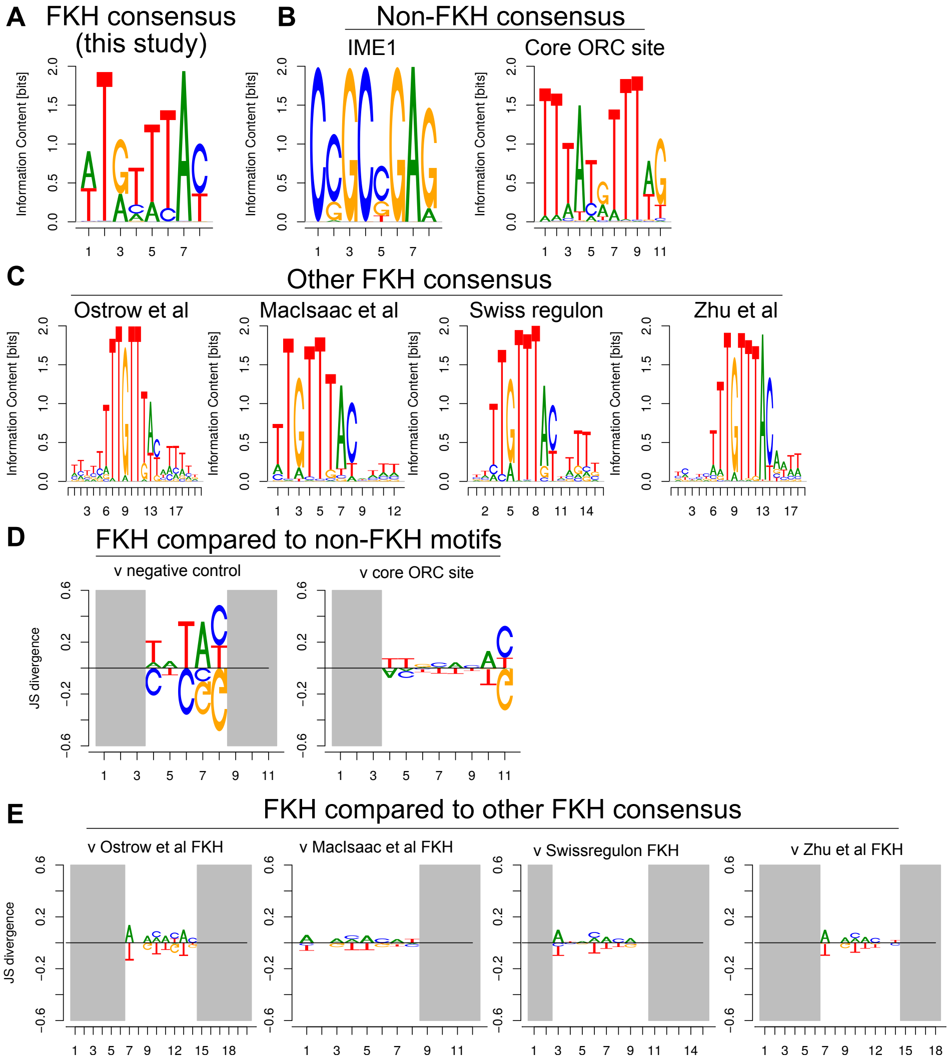


**Figure S4.** Challenging FKH consensus motif from the Fkh1/2 ChIPchip. **A)** Sequence logo of the position weight matrix of the FKH consensus site in this study used to map motifs in Figure 4B. **B)** Sequence logos of two non-FKH position weight matrixes. Left, IME1, represents a motif distinct from FKH. Right, core ORC site, represents a motif similar to that of FKH1. **C)** Sequence logos of previously annotated FKH1 consensus. **D)** Comparison of the position weight matrices from this study's FKH consensus to that of two control motifs, a negative control from a distinct motif with a visually distinct consensus (IME1) and the ORC site consensus. **E)** Comparison of the position weight matrices from this study's FKH consensus to that of FKH consensus published in previous studies [11–14].

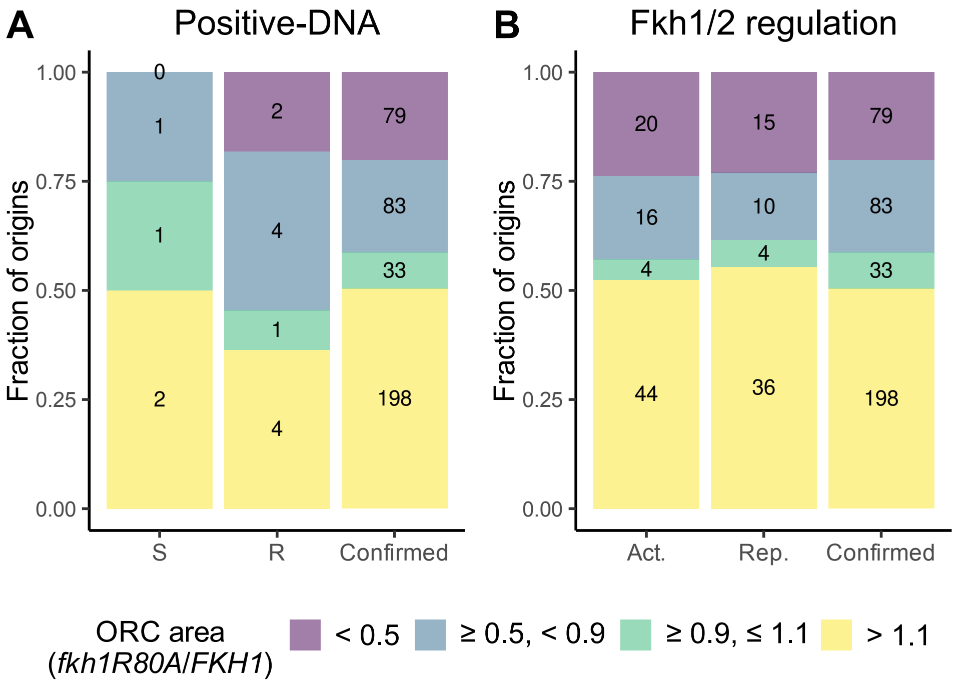


Figure S5. Distributon of fkh1R80A/FKH1 ORC binding ratios as descscribed in Figure 6 for A) FHA-dependent (n = 4, S) and FHA-independent (n = 11, R) positive-DNA origins and B) Fkh1/2-activated (Act., n = 84)) and repressed (Rep., n = 65) origins. There were 393 confirmed yeast chromosomal origins used for this study. The *fkh1R80A/FKH1* ORC binding ratio was determined by dividing the the total sum of ORC binding signal areas in *fkh1R80A* cells by the area in *FKH1* cells spanning nucleotides -100 to +100 of each origin. The origins were parsed by their *fkh1R80A/FKH1* ORC binding ratios into the four categories indicated: 1, purple origins showed < 50% the ORC binding signal in *fkh1R80A* compared to *FKH1* cells; 2, blue origins showed at least a 10% reduction but nor more than a 50% reduction; 3, green origins were unaffected; 4,yellow origins showed greater than a 10% enhanced ORC binding in *fkh1R80A* relative to *FKH1* cells. Enrichments were challenged with the Hypergeometric Distribution function comparing cohort ratios to that measured within all confirmed origins. Origin cohorts in A were defined in this study. Origin cohorts in B were defined in [10].

#### Supplemental References


1. Shor, E. *et al.* The origin recognition complex interacts with a subset of metabolic genes tightly linked to origins of replication. *PLoS Genet* **5,** e1000755 (Dec. 2009).
2. Casey, L., Patterson, E., Müller, U. & Fox, C. Conversion of a replication origin to a silencer through a pathway shared by a Forkhead transcription factor and an S phase cyclin. *Mol Biol Cell* **19,** 608–22 (Feb. 2008).
3. Dummer, A. *et al.* Binding of the Fkh1 Forkhead Associated Domain to a Phosphopeptide within the Mph1 DNA Helicase Regulates Mating-Type Switching in Budding Yeast. *PLoS Genet* **12,** e1006094 (June 2016).
4. Skene, P. & Henikoff, S. A simple method for generating high-resolution maps of genome-wide protein binding. *Elife* **4,** e09225 (June 2015).
5. Buck, M., Nobel, A. & Lieb, J. ChIPOTle: a user-friendly tool for the analysis of ChIP-chip data. *Genome Biol* **6,** R97 (2005).
6. Homann, O. & Johnson, A. MochiView: versatile software for genome browsing and DNA motif analysis. *BMC Biol* **8,** 49 (Apr. 2010).
7. Nettling, M. *et al.* DiffLogo: a comparative visualization of sequence motifs. *BMC Bioinformatics* **16,** 387 (Nov. 2015).
8. Feng, W. *et al.* Genomic mapping of single-stranded DNA in hydroxyurea-challenged yeasts identifies origins of replication. *Nat Cell Biol* **8,** 148–55 (Feb. 2006).
9. Crabbé, L. *et al.* Analysis of replication profiles reveals key role of RFC-Ctf18 in yeast replication stress response. *Nat Struct Mol Biol* **17,** 1391–7 (Nov. 2010).
10. Knott, S. *et al.* Forkhead transcription factors establish origin timing and long-range clustering in S. cerevisiae. *Cell* **148,** 99–111 (Jan. 2012).
11. MacIsaac, K. *et al.* An improved map of conserved regulatory sites for Saccharomyces cerevisiae. *BMC Bioinformatics* **7,** 113 (Mar. 2006).
12. Pachkov, M., Balwierz, P., Arnold, P., Ozonov, E. & van, N. E. SwissRegulon, a database of genome-wide annotations of regulatory sites: recent updates. *Nucleic Acids Res* **41,** D214–20 (Jan. 2013).
13. Zhu, C. *et al.* High-resolution DNA-binding specificity analysis of yeast transcription factors. *Genome Res* **19,** 556–66 (Apr. 2009).
14. Ostrow, A. *et al.* Fkh1 and Fkh2 bind multiple chromosomal elements in the S. cerevisiae genome with distinct specificities and cell cycle dynamics. *PLoS One* **9,** e87647 (2014).
